# Molecular and cognitive signatures of ageing partially restored through synthetic delivery of IL2 to the brain

**DOI:** 10.1101/2022.03.01.482519

**Authors:** Pierre Lemaitre, Samar Tareen, Emanuela Pasciuto, Loriana Mascali, Araks Martirosyan, Zsuzsanna Callaerts-Vegh, James Dooley, Matthew G. Holt, Lidia Yshii, Adrian Liston

## Abstract

Cognitive decline is a common pathological outcome during aging, with an ill-defined cellular or molecular basis. Among the cellular changes observed with age are alterations to neuronal plasticity, changes in the glial compartment and the decline of the neurogenic niche. In the recent years, the concept of inflammaging, defined as a low-grade inflammation increasing with age, has emerged as a nexus for age-related diseases. This increase of basal inflammation is also observed in the central nervous system. While not classically considered a neurological cell type, infiltrating T cells increase in the brain with age, and may be responsible for amplification of inflammatory cascades and disruptions to the neurogenic niche. Recently, a small resident population of regulatory T cells has been identified in the brain, and the capacity of IL2-mediated expansion of this population to counter neuroinflammatory disease has been demonstrated. Here we test a brain-specific IL2 delivery system for the prevention of neurological decline in aging mice. We identify the molecular hallmarks of aging in the brain glial compartments, and identify partial restoration of this signature through IL2 treatment. At a behavioral level, brain IL2 delivery prevented the age-induced defect in spatial learning, without improving the general decline in motor skill or arousal. These results identify immune modulation as a potential path to preserving cognitive function for healthy ageing.

## Introduction

Aging is an irreversible process associated with physical deterioration and hence especially noticeable in organs containing mainly post-mitotic cells, such as the brain. The progressive decline in memory, orientation, attention, motivation and cognition seen in humans is thought to stem largely from aging-related processes [1]. Although cognitive aging research generally focuses on neurons, which are widely held as the computational units of the brain, there is an increasingly realization that reciprocal interactions between neurons and non-neuronal cells (collectively referred to as glia) play key roles in all aspects of nervous system function. These include the regulation of synapse formation and function by astrocytes, as well as myelination and provision of trophic support to neurons by oligodendrocytes. Once thought of as solely immune cells, which protect the CNS against injury and disease, there is increasing evidence that the same systems used by microglia to remove invading pathogens from the brain have also been co-opted to remove weak and/or inappropriate neuronal synapses, establishing precise patterns of neuronal connectivity in the brain [2]. It should not be surprising, therefore, that aging-related cognitive decline has been associated with several molecular processes, including chronic inflammation, impaired autophagy, macromolecular damage, and mitochondrial dysfunction and senescence [3], which impact the function of all CNS cell types [4–7].

In addition to the shared cellular phenotypes of ageing, unique phenotypes emerge among the individual glial cell types with ageing. Microglia from aged animals are hyper-reactive to inflammatory stimuli, with higher basal levels of cytokine expression and exaggerated responses to activation [8]. Aged microglia exhibit shortening of processes, slower process movement, and enlarged soma volumes [9]. Functional decline of microglia is hypothesized to be progressive over the lifespan, due to accumulation of oxidative DNA damage and non-degradable protein and lipid aggregates. This inflamed and dysfunctional microglial profile of old age plays a role in the development of age-associated neurodegenerative diseases [10]. The ageing phenotype of microglia is driven in part by the breakdown of myelin [11], a phenomenon paralleled by the ageing of oligodendrocytes. Oligodendrocytes are derived from specific neural progenitor cells, oligodendrocyte progenitor cells (OPCs), with the primary role of myelin production. OPC heterogeneity increases with age [12], while the myelination function of oligodendrocytes is decreased [13], with an associated breakdown in myelin integrity [14]. Astrocytes undergo substantial morphological and transcriptional changes with ageing, consistent with the acquisition of a reactive pro-inflammatory phenotype [15, 16], although there are regional differences in the response of astrocytes across different areas of the brain [17]. The degree to which these glial changes are drivers, mediators or bystanders in the cellular pathways linking ageing to cognitive decline is debated.

One of the most recognized effects of aging is dysregulation of the immune system. Both immunosenescence, defective initiation and resolution of immune responses, and inflammaging occur with age [18, 19]. Aging affects both the adaptive and the innate immune system [20, 21]. In the brain, immune aging is accompanied by a low-grade chronic proinflammatory environment, characterized by increased production of proinflammatory cytokines, such as interleukin-6 (IL-6) and tumor necrosis factor alpha (TNFα), acute-phase proteins, reactive oxygen species (ROS), and autoantibodies. Lymphatic drainage of inflammatory products from the brain becomes insufficient with age, contributing to cognitive decline [22]. CNS leukocytes may also play a role in age-related inflammation and neurodegeneration [23–25]. In particular, the accumulation of clonally expanded T cells is observed in the aged brain, accompanied by production of IFNγ [26] and loss of CCR7 [27]. The inflammatory effects of T cells in the brain in turn impede the neuronal stem cell niche and alter the biology of the glial compartment, amplifying the effects of neuroageing [26].

Beyond the observed age-associated cognitive decline, ageing is also a major risk factor for pathological neurodegeneration. The risk of neurodegenerative diseases such as Alzheimer’s disease (AD), Parkinson’s disease (PD), amyotrophic lateral sclerosis (ALS), and primary progressive multiple sclerosis (PPMS) increases significantly in the elderly population [3]. This enhanced rate of neurodegenerative disease is associated with neuroinflammation, characterized by reactive central nervous system microglia and astroglia, as well as infiltrating peripheral monocytes and lymphocytes. The presence of vascular and parenchymal T cells in brains of postmortem AD patients [28] and in transgenic models of AD [29], suggests the potential involvement of T cells in the development of neuroinflammation. In PD, T cells have been observed in the substantia nigra of patients, and pharmacological manipulation of T cell responses alters disease outcome in mouse models [30]. Likewise, T cell infiltration is observed at sites of motor neuron loss in ALS patients, with modified disease progression in T cell-deficient mouse models [31]. The association of HLA alleles to AD and PD susceptibility supports a causative role of T cells in the process [32–34]. Together, this suggests that infiltrating T cells in the brain contribute to both the common age-related cognitive decline and a diversity of neurodegenerative diseases.

Regulatory T cells (Treg) are a subset of T cells with potent anti-inflammatory and pro-repair functions. Treg reside in both lymphoid and in non-lymphoid organs, where they exert diverse functions regulating tissue homeostasis and contributing to tissue repair [35]. Tregs are found in low numbers in the mouse and human brain, and are observed in both the parenchyma and meningeal lymphatics [36]. Tregs have been proposed to have a protective function in a variety of neuroinflammatory and/or neurodegenerative diseases, with protective effects observed in mouse models of MS, AD, PD, ALS and stroke, among others [37–41]. Proposed therapeutic strategies to harness Tregs are typically based on either direct delivery of Tregs, through cell-based approaches, or the provision of IL2, the key survival factor for Tregs [42]. Despite the growing recognition of a protective role of IL2 and Tregs in neuroinflammatory and neurodegenerative processes, the potential of these cells to protect the ageing brain remain unknown. Here, we used the synthetic delivery of brain-specific IL2, which was recently demonstrated to protect against neuroinflammatory injury and disease [43], to test the hypothesis that neurological ageing can be restrained by modulation of the immunological compartment of the brain. Using young and aged mice we characterized the molecular signature of aging in the brain stroma, and found IL2 capable of restraining a core component of the molecular aging process across glial types. These molecular effects were associated with a partial preservation of cognitive capacity, suggesting a potential use of brain-specific IL2 production to combat age-related cognitive decline.

## Results

### Age drives a change in the cellular composition of brain-resident glia

A key limitation to proposed Treg- or IL2-based therapies is the peripheral impact of most proposed strategies, with the potential for undesirable peripheral immunosuppression. The use of brain-targeted delivery opens the door to potential utilization for neuropathologies, while preserving the integrity of the peripheral immune system. One way to achieve brain-specific expression of IL2 is through the gene-delivery vector *PHP.GFAP*-IL2. This approach results in the restricted production of IL2 in the brain, and an accompanying expansion of brain-resident Tregs [43]. PHP.*GFAP*-IL2 treatment is protective against neuroinflammation in the context of traumatic brain injury, stroke and experimental autoimmune encephalitis [43]. In order to test the influence of brain-specific IL2 delivery on ageing, we treated young mice, at 2 months of age, and old mice, at 22 months of age, with *PHP.GFAP*-IL2 or the control PHP.*GFAP*-GFP vector. The effect of this treatment was assessed on the glial compartment of the whole brain two months later, at 4 and 24 months of age, through the flow cytometric sorting and single cell RNA sequencing of microglia, astrocytes and oligodendrocytes (**Figure 1A**). The target glial populations (**Supplementary Figure 1**), and minor population contaminants (**Supplementary Figure 2**) were identified through expression analysis of key lineage markers and integration with reference datasets [44]. PCA analysis of major and minor population frequencies found an association with age but not treatment, indicating an age-dependent phenotype (**Figure 1B**). Reclustering of microglia resulted in tightly clustered cells (**Figure 1C**), while reclustering of oligodendrocytes led to a clear separation of oligodendrocyte precursors (OPCs) and oligodendrocytes (**Figure 1D**). Reclustering of astrocytes, by contrast, produced a clear transcriptional separation of Bergmann glia, Cerebellar astrocytes, Olfactory astrocytes, striatal astrocytes, telencephalon astrocytes and non-telencephalon astrocytes (**Figure 1E**), based on key markers and data-set integration (**Supplementary Figure 3**).

**Figure 1.**
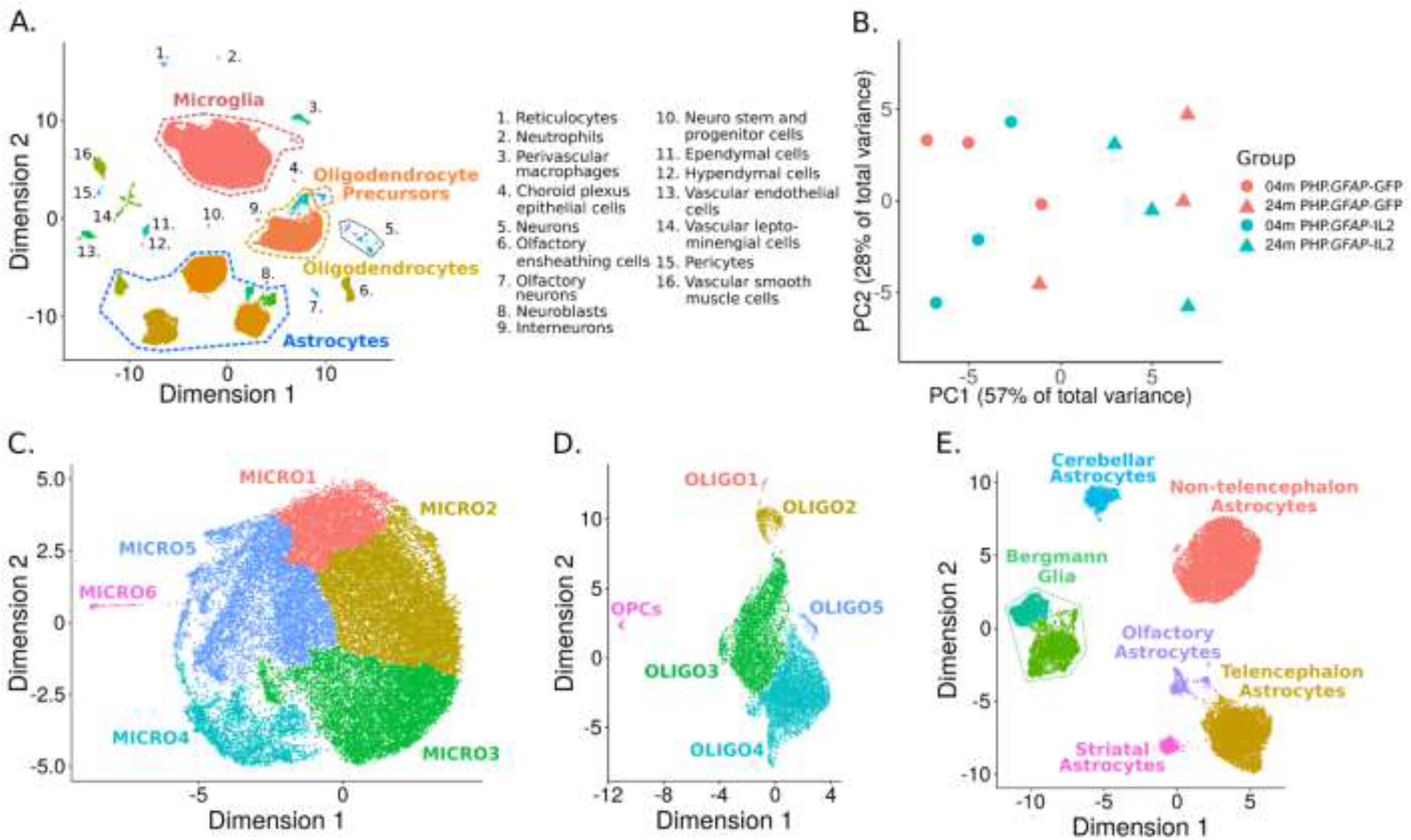
Single cell sequencing identifies a shift in glia proportions with age. Wildtype mice were treated with PHP. *GFAP*-IL2 (or PHP. *GFAP*-GFP control vector) at 2 months or 22 months of age (n=3/group). Two months post-treatment, the glial compartment was sorted from perfused mice and assessed using 10x single-cell transcriptomics. **A)** UMAP projection of the cells from the dataset. Colored names indicate the glial populations sorted for downstream analysis (as in Figure S1), whereas minor populations annotated in black are contaminant cell types found in the data post-sorting (as in Figure S2). **B)**PCA was performed on the proportions of cells from each sample in each cluster to generate an overview of whether age and/or treatment show any global trend with respect to sample proportions in each cluster of the data. **C)** The microglia clusters were reclustered and projected in UMAP to generate further separation in the data. **D)** The oligodendrocyte and oligodendrocyte precursor (OPC) clusters were reclustered and projected in UMAP to generate further separation in the data. **E)** The astrocyte clusters were reclustered and reprojected in UMAP. The different astrocyte clusters were then mapped onto known astrocyte subtypes.

We first investigated cellular changes in the microglial compartment. Reclustered microglia were assessed for separation based on age and treatment (**Figure 2A,B**). Visual separation of age was observed on the UMAP, with quantification of clusters identifying increased representation of Micro1 and Micro2 clusters in young mice, with Micro3 to Micro6 clusters increased in aged mice (**Figure 2C**). Based on the prior characterization of disease-associated microglia (DAM), we identified microglia expressing characteristic markers (**Supplementary Figure 4**) and classified cells as DAM based on aggregate marker expression above a threshold (**Figure 2D**) calibrated against published databases [45]. DAM were nearly absent in young mice, with a significant increase in aged mice (**Figure 2E**), on the order of ~2% of microglia. For each identified subset, no significant changes were observed due to IL2 treatment (**Figure 2C,E**). This data suggests that brain-specific IL2 supplementation does not alter the microglial progression through cellular states normally observed with age.

**Figure 2.**
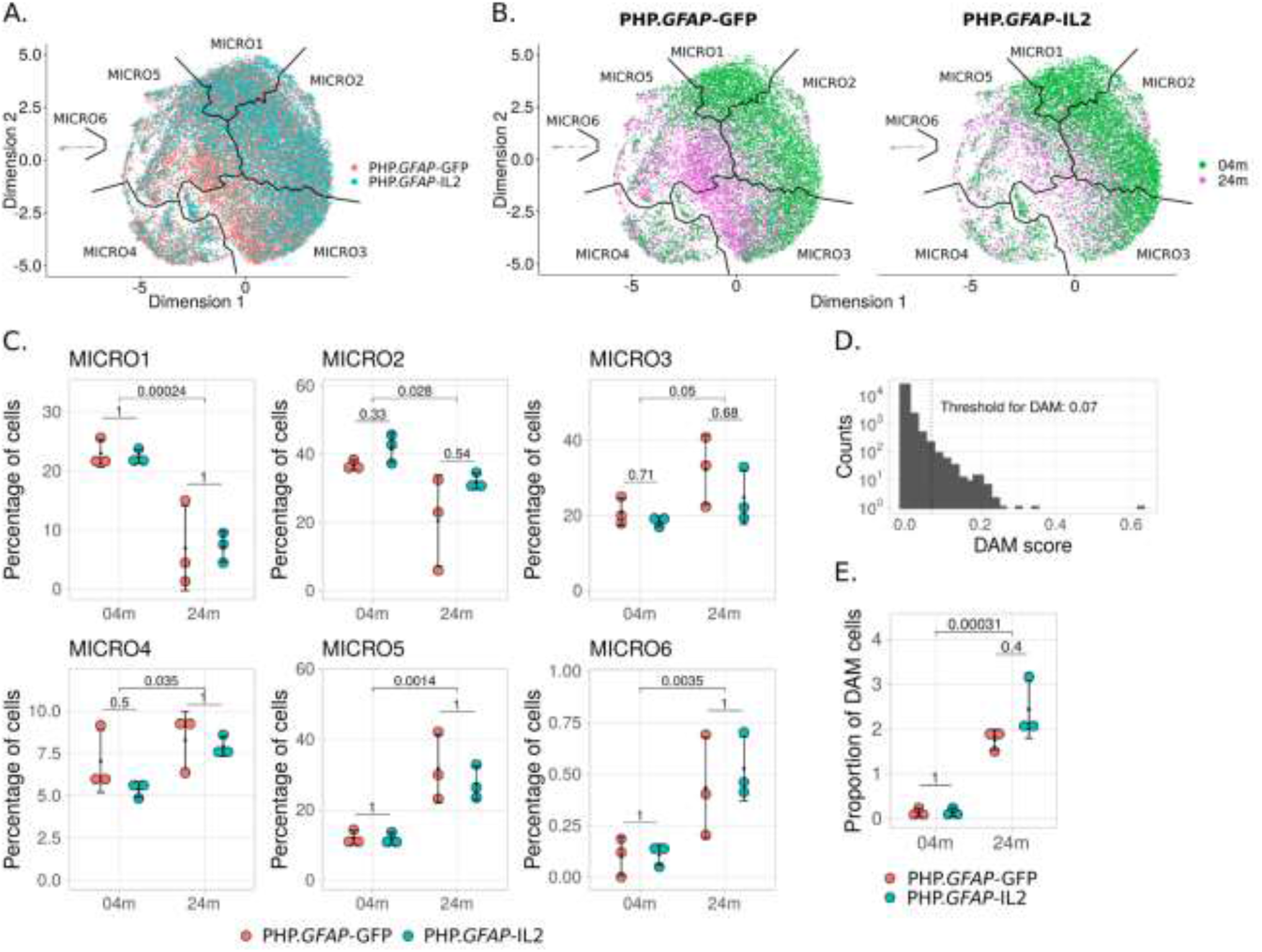
Age-dependent accumulation of disease-associated microglia is intact in IL2-treated mice. Wildtype mice were treated with PHP.*GFAP*-IL2 (or PHP.*GFAP*-GFP control vector) at 2 months or 22 months of age (n=3/group). Two months post-treatment, the glial compartment was sorted from perfused mice and assessed using 10x single-cell transcriptomics. Microglia were reclustered and projected in UMAP. **A)** Distribution of the microglia by treatment. **B)** Distribution of microglia by age, with subpanels separating the cells by treatment (PHP.*GFAP*-IL2 or PHP.*GFAP*-GFP control vector). **C)** Proportions of cells from each sample, contained within each of six microglia clusters. Each panel shows multiple-testing corrected p-values for t-tests comparing the means of the samples between treatments within each age group, and the means of the two age groups irrespective of treatment. **D)** Histogram of disease-associated microglia (DAM) scoring for microglia, indicating the 0.07 threshold for classification as DAM. **E)** Proportions of DAM cells in each sample, with multiple-testing corrected p-values for t-tests comparing the means of the samples between treatments within each age group, and the means of the two age groups irrespective of treatment.

Reclustering of oligodendrocytes resulted in a clear separation of OPCs, with the remaining oligodendrocyte population showing a visual separation based on age (**Figure 3A,B**). While OPC numbers were unaffected by ageing, the clusters Oligo1 and Oligo2 were reduced with age (**Figure 3C**). Pseudotime analysis allowed the generation of branching trajectory trees across each sample (**Supplementary Figure 5**). Compared to the young IL2-treated mice, aged mice demonstrated altered proportions of cells in different stages of the pseudo-time trajectory, with substantial deviations in the relative distribution of cells (**Figure 3D**). In concordance, pseudotime reconstruction of oligodendrocyte progression identified Oligo1, Oligo2 and part of Oligo3 clusters as the most immature state, with progression through Oligo3, Oligo4 and Oligo5 with maturation (**Figure 3E, Supplementary Figure 5**). This reconstruction aligned with the expression of S100β (**Figure 3F**), which, in oligodendrocytes, serves as a canonical maturation marker, and other oligodendrocyte markers (**Supplementary Figure 6**). Together, these results suggest a structured progression from immature to mature oligodendrocytes occurring in aged mice. As with microglia, brain-specific delivery of IL2 through *PHP.GFAP*-IL2 did not alter this normal maturational progression (**Figure 3C,D**).

**Figure 3.**
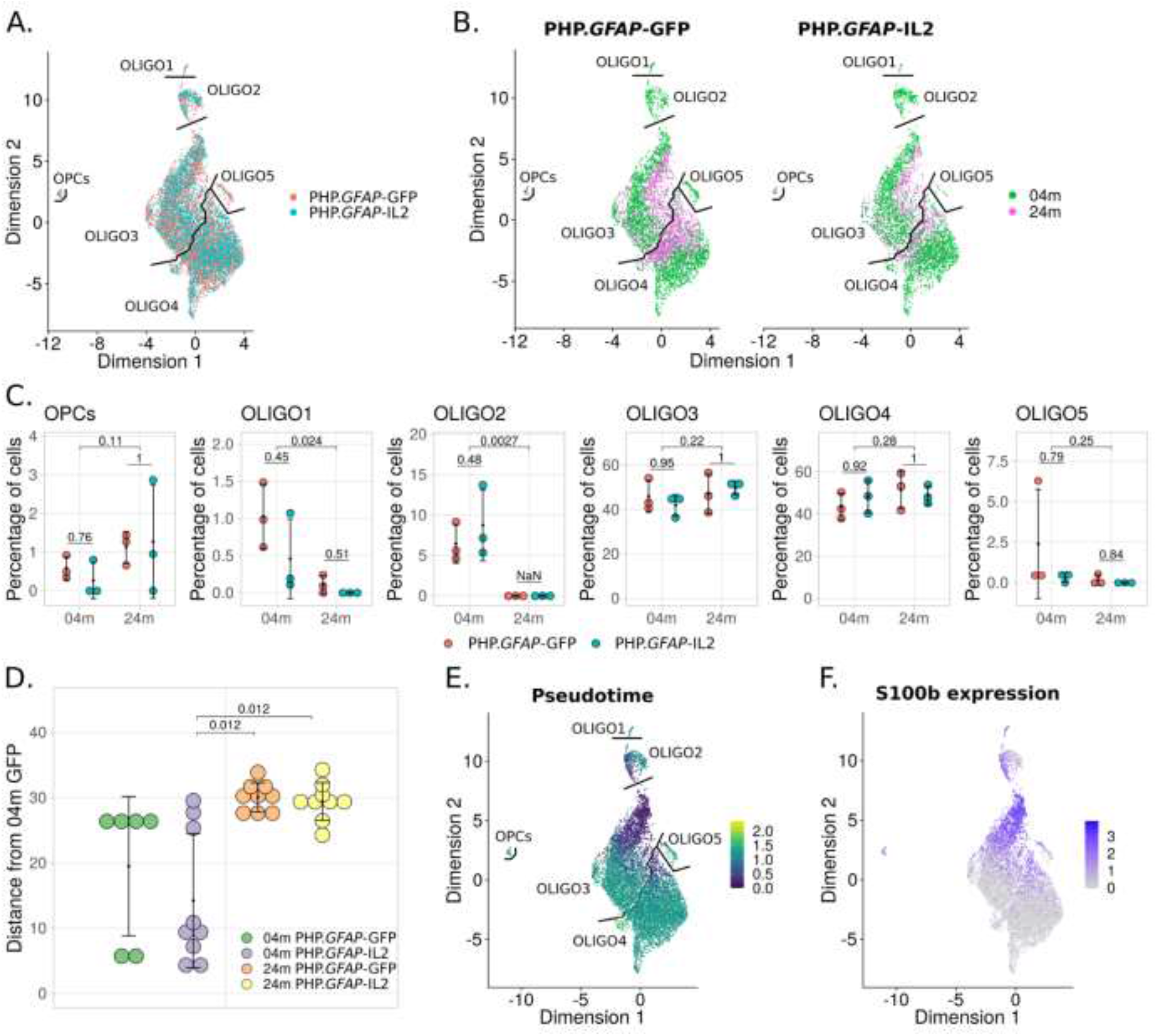
Normal age-dependent cellular progression for oligodendrocytes following IL2-treatment. Wildtype mice were treated with PHP.*GFAP*-IL2 (or PHP.*GFAP*-GFP control vector) at 2 months or 22 months of age (n=3/group). Two months post-treatment, the glial compartment was sorted from perfused mice and assessed using 10x single-cell transcriptomics. Oligodendrocytes and OPC clusters were reclustered and projected in UMAP space. **A)** Distribution of the oligodendrocytes and precursors by treatment. **B)** Distribution of the oligodendrocytes and precursors by sample age, with subpanels separating the cells by treatment. **C)** Proportions of cells in the various oligodendrocyte and OPC clusters. Each panel shows multiple-testing corrected p-values for t-tests comparing the means of the samples between treatments within each age group, and the means of the two age groups irrespective of treatment. **D)** The sum of deviance plot summarizing the proportions of age groups in the trajectory trees generated for oligodendrocytes and precursors (Figure S5). The plot visualizes the difference of proportion of cells by age in each branch of the respective trajectory tree relative to young PHP.*GFAP*-GFP treated mice, illustrating which age and treatment group shows the most deviation in the cell trajectory relative to the reference. **E)** Pseudotime projection calculated using the DDRTree algorithm in Monocle v2 mapped onto the UMAP space. **F)** *S100β* expression superimposed on the UMAP projection.

Finally, we assessed the effect of ageing and IL2 treatment on astrocyte maturation (**Figure 4A**). Unlike microglia and oligodendrocytes, astrocytes were highly heterogeneous, corresponding to previous anatomical separation of subsets [44, 46]. Visual inspection of UMAP reclustering found an age-based segregation within clusters (**Figure 4B**). At the subtype level, only minor shifts were observed with age – a decrease in Bergmann glia and striatal astrocytes, and an increase in non-telencephalon astrocytes (**Figure 4C**). To investigate the effect of ageing within each subtype, we constructed pseudotime maturational trees (**Supplementary Figure 7**). To assess the effect of age and treatment on maturation, we generated a summary statistic of the sum of frequency difference in each tree branch. Overall, ageing in control-treated groups (PHP.*GFAP*-GFP) drove a trend towards aggregate change in Bergmann glia, cerebellar astrocytes and striatal astrocytes (**Figure 4D**). Treatment with PHP.*GFAP*-IL2 did not alter the trajectory progression in young mice, however it partially prevented the aged trajectory distribution developing in cerebellar astrocytes (**Figure 4D**). Together, this analysis found substantial age-induced transcriptome changes in all major glial cell types, with no transcriptional abnormalities caused by IL2-delivery.

**Figure 4.**
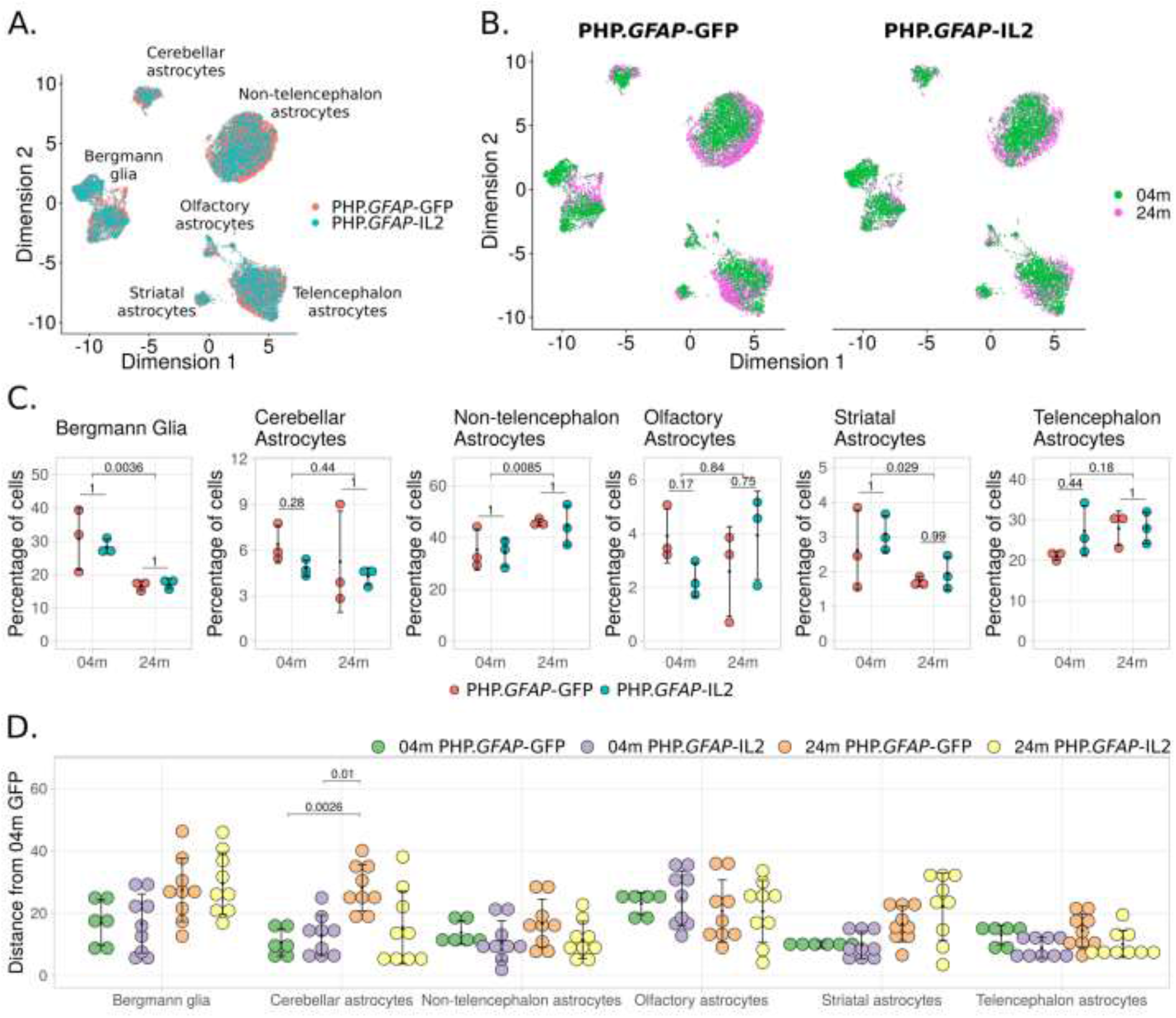
Partial prevention of age-dependent transcriptional trajectory in astrocytes following IL2-treatment. Wildtype mice were treated with *PHP.GFAP*-IL2 (or PHP.*GFAP*-GFP control vector) at 2 months or 22 months of age (n=3/group). Two months posttreatment, the glial compartment was sorted from perfused mice and assessed using 10x single-cell transcriptomics. Astrocyte clusters were reclustered and reprojected in UMAP space, with subset annotation based on key marker expression and mapping onto known astrocyte subtypes. **A)** Distribution of the different astrocyte subtypes by treatment. **B)** Distribution of the different astrocyte subtypes by age, with subpanels separating the cells by treatment. **C)** Proportions of cells from each group that fall into each astrocyte subtype. Each panel shows multiple-testing corrected p-values for t-tests comparing the means of the samples between treatments within each age group, and the means of the two age groups irrespective of treatment. **D)** The sum of deviance plot summarizing deviations in the proportion of cells in each branch of the pseudotime trajectory, compared to that of the young control, generated for each astrocyte subtype (Figure S7). The plot visualizes the difference of proportion of cells by age in each branch of the respective trajectory tree relative to young PHP.*GF4P*-GFP treated mice, illustrating which age and treatment group shows the most deviation in the cell trajectory relative to the reference.

### Brain-specific IL2 delivery mitigates age-induced molecular changes in resident glia

Having established that local IL2 expression in the brain does not distort the relative frequency of major glial cell types, nor the normal trajectory distribution, we sought to investigate the impact on age-induced molecular pathways. First we performed a differential expression analysis for each major glial cell type (**Supplementary Resource 1**). Twodimensional differential expression comparisons demonstrated that the largest transcriptional changes were occurring with age, for each glial cell type (**Figure 5**). The change occurring with treatment of aged mice was reduced in scale, but ran directly counter to the age-induced changes. Correlation of comparative differential expression indicated that local IL2 treatment countered 47%, 55% and 46% of the age-induced transcriptional changes in microglia, oligodendrocytes and astrocytes, respectively (**Figure 5**). We next sought to determine the effect at the gene set level, where accumulated minor changes in a molecular pathway can be assessed. Gene set enrichment was performed on each glial cell type, comparing the effects of age and treatment. Among the key pathways modified by ageing in microglia were those of proteostasis-related pathways, autophagy-associated pathways, key signaling pathways (including Ras, PI3K and mTOR pathways), neurodegeneration-associated genes and the cellular senescence pathway (**Supplementary Resource 2**). Oligodendrocytes and astrocytes underwent a similar transcriptional change with age, albeit with fewer pathways significantly altered, with altered expression of proteostasis pathways, autophagy-associated pathways and neurodegeneration-associated genes (**Supplementary Resource 2**). To assess the effect of treating aged mice with local IL2 production, we created a directionality map, where each transcriptional change within a gene set was compared to the scale and direction of the change occurring with age in the control mouse. This allowed the direct comparison of transcriptional changes occurring within each pathway, across each of the glial cell types. Almost all key age-induced pathway changes were also mirrored in IL2-treated mice, with 26/26 pathways in microglia, 10/11 pathways in oligodendrocytes and 15/16 pathways in astrocytes undergoing the same direction of transcriptional change in aged IL2-treated mice as occurred in aged control mice (**Figure 6**). However in almost all cases (26/26 pathways in microglia, 10/11 pathways in oligodendrocytes and 16/16 pathways in astrocytes), IL2-treatment of aged mice reverted the expression profile to that of the young control mouse (**Figure 6**). Together, these results demonstrate that the key coordinated transcriptional changes occurring in aged glial cells, in protein production pathways, autophagy-associated pathways and neurodegeneration-associated genes, are partially reverted through a two month treatment regime of locally-produced IL2.

**Figure 5.**
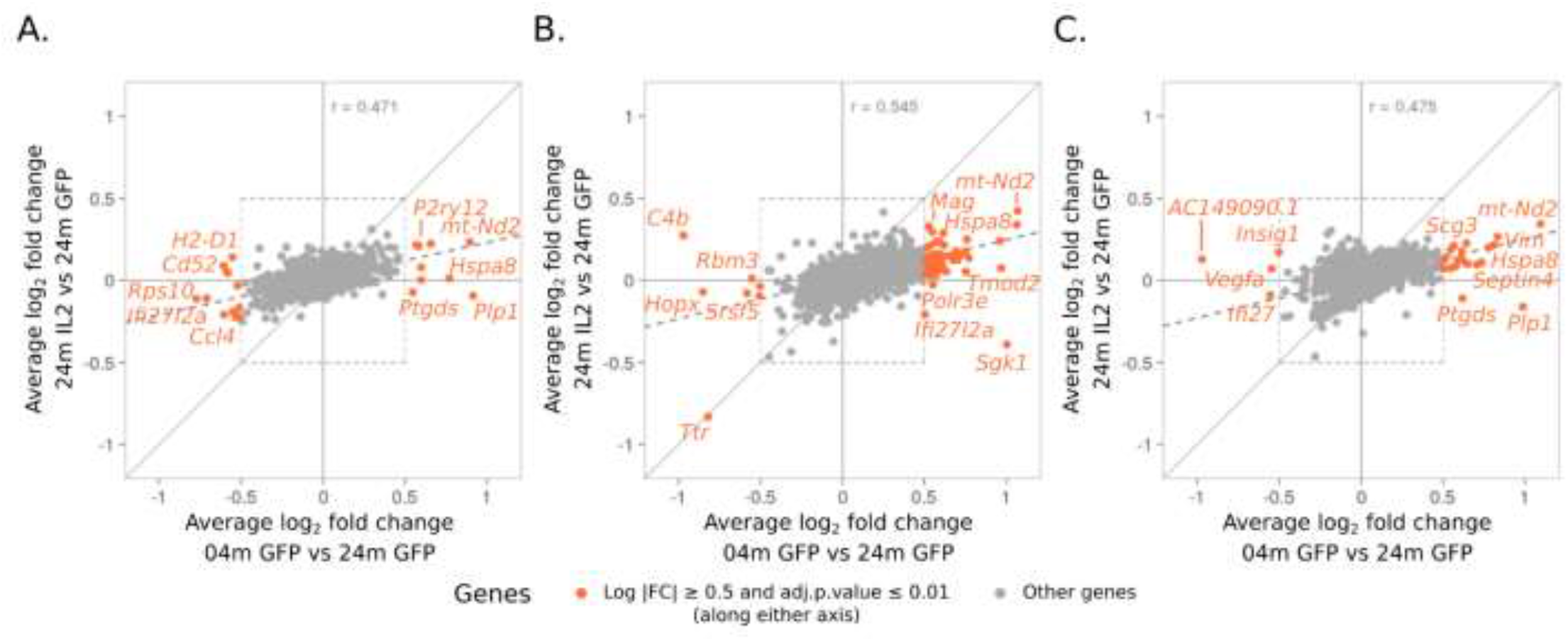
Local IL2 production in aged mice partially reverts the molecular changes occurring in aged glia. Two-dimensional differential expression plots illustrating the contrast for normal aging (young PHP.*GFAP*-GFP mice versus aged PHP.*GFAP*-GFP mice) to the contrast for treatment-induced changes in aged mice (aged PHP.*GFAP*-IL2 mice versus aged PHP.*GFAP*-GFP). Expression plots shown for **A)** microglia, **B)** oligodendrocytes and OPCs, and **C)** astrocytes (total).

**Figure 6.**
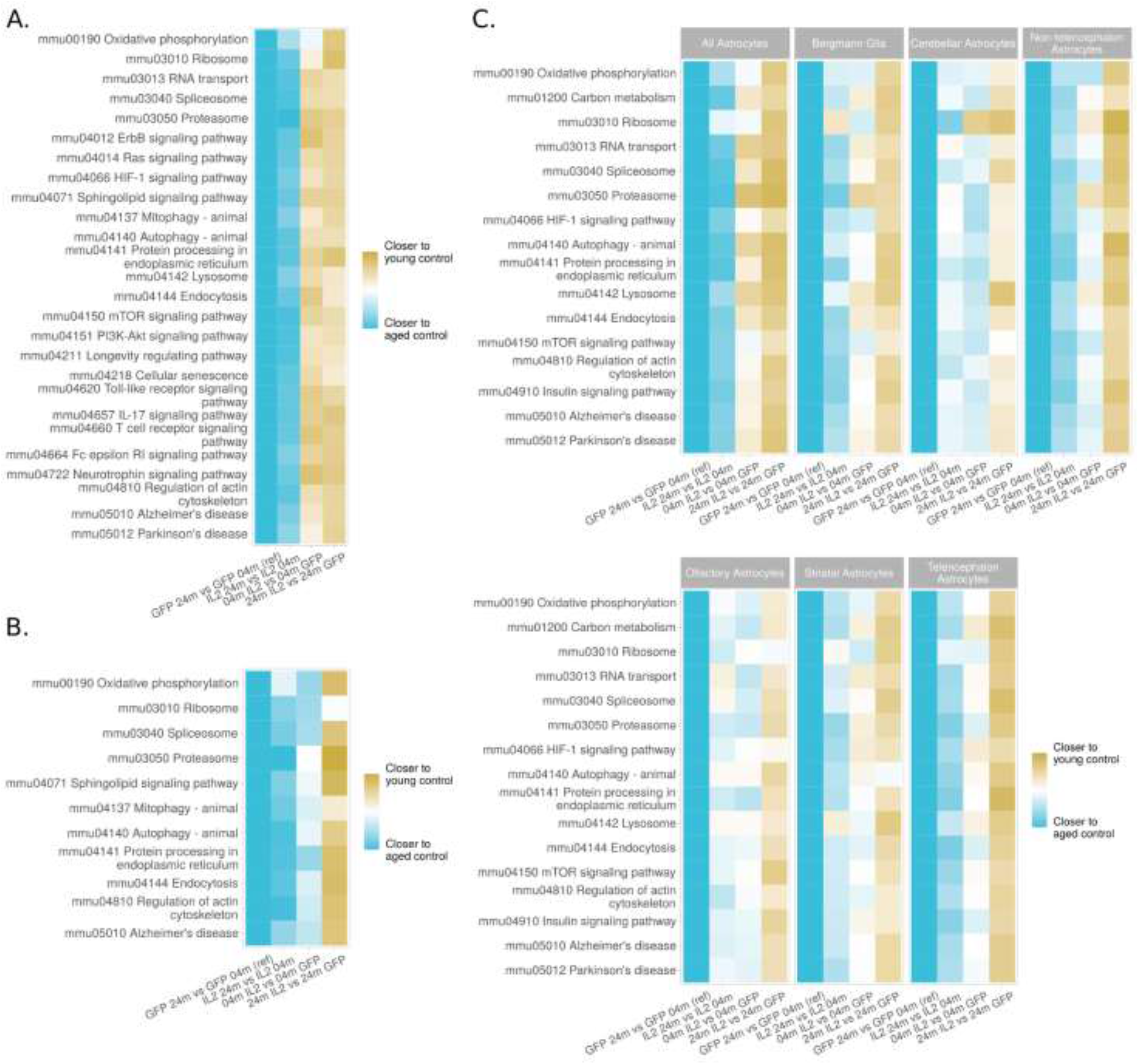
Age-induced pathways in glia are reverted to the young transcriptional state by local IL2 production. Gene set enrichment was performed based on differential expression across the comparisons for age (aged PHP.*GFAP*-GFP mice versus young PHP.*GFAP*-GFP mice), treatment in young mice (young PHP.*GFAP*-GFP mice versus young PHP.*GFAP*-IL2 mice), treatment in old mice (aged PHP.*GFAP*-IL2 mice versus aged PHP.*GFAP*-GFP mice) and age in treated mice (aged PHP.*GFAP*-IL2 mice versus young PHP.*GFAP*-IL2 mice). Gene sets were taken from the GAGE library and curated to the respective KEGG pathway. Listed pathways were manually curated for relevance to glial biology and non-redundancy from the unbiased list enriched in at least one of the four respective differential expression contrasts. For each pathway, directionality maps were generated, summarizing the change in direction of gene expression for member genes relative to the expression change with age (aged PHP.*GFAP*-GFP mice contrasted with young PHP.*GFAP*-GFP mice). Directionality is shown for each pathway across each of the three contrasts for **A)** microglia, **B)** oligodendrocytes and OPCs, and **C)** total astrocytes and the identified astrocyte subsets.

### Brain-specific IL2 gene-delivery provides partial protection for cognitive decline

Based on the ability of PHP.*GFAP*-IL2 to partially prevent the key coordinated transcriptional signature of ageing in glia, we sought to determine whether these molecular changes would result in improvements at the cognitive level. We treated old mice with PHP.*GFAP*-IL2 or control PHP.*GFAP*-GFP vector, and initiated a battery of behavioral tests two months later, at 24 months of age, compared to a 4 month-old control-treatment group. Aged mice demonstrated multiple signs of behavioral decline, with reduced activity in their home cage (**Figure 7A**), impaired mobility on the rotarod (**Figure 7B**) and reduced exploration in an open field test (**Figure 7C**). Aged mice also demonstrated decline in the social novelty-seeking behavior of the sociability test (**Figure 7D**) and reduced explorative behavior in the light-dark test (**Figure 7E**). For each of these measures, which may reflect a reduction in physical capacity and decreased arousal, brain-specific IL2 delivery did not alter the age-related decline. Finally, we tested spatial learning in in the Morris water maze. Aged mice demonstrated poor performance in the Morris water maze in two aspects. First, aged mice had a large reduction in swim velocity during the test (**Figure 7F**), a phenotype likely to be derived from physical rather than cognitive decline. Second, aged mice showed delayed spatial learning acquisition during training (**Figure 7G-I**) and demonstrated a reduced preference for the target quadrant after 10 days of training (**Figure 7H**), a phenotype reflecting reduced cognitive capacity for spatial memory formation. Treatment of aged mice with PHP.*GFAP*-IL2 had no impact on the physical decline in swim velocity, but was able to completely correct the defective observed in spatial memory formation (**Figure 7F,H**). Together, these results suggest that PHP.*GFAP*-IL2, even when delivered to old mice, is able to prevent or restore cognitive decline in spatial memory formation, without impacting other aspects of behavioral age-related decline, such as physical prowess and arousal.

**Figure 7.**
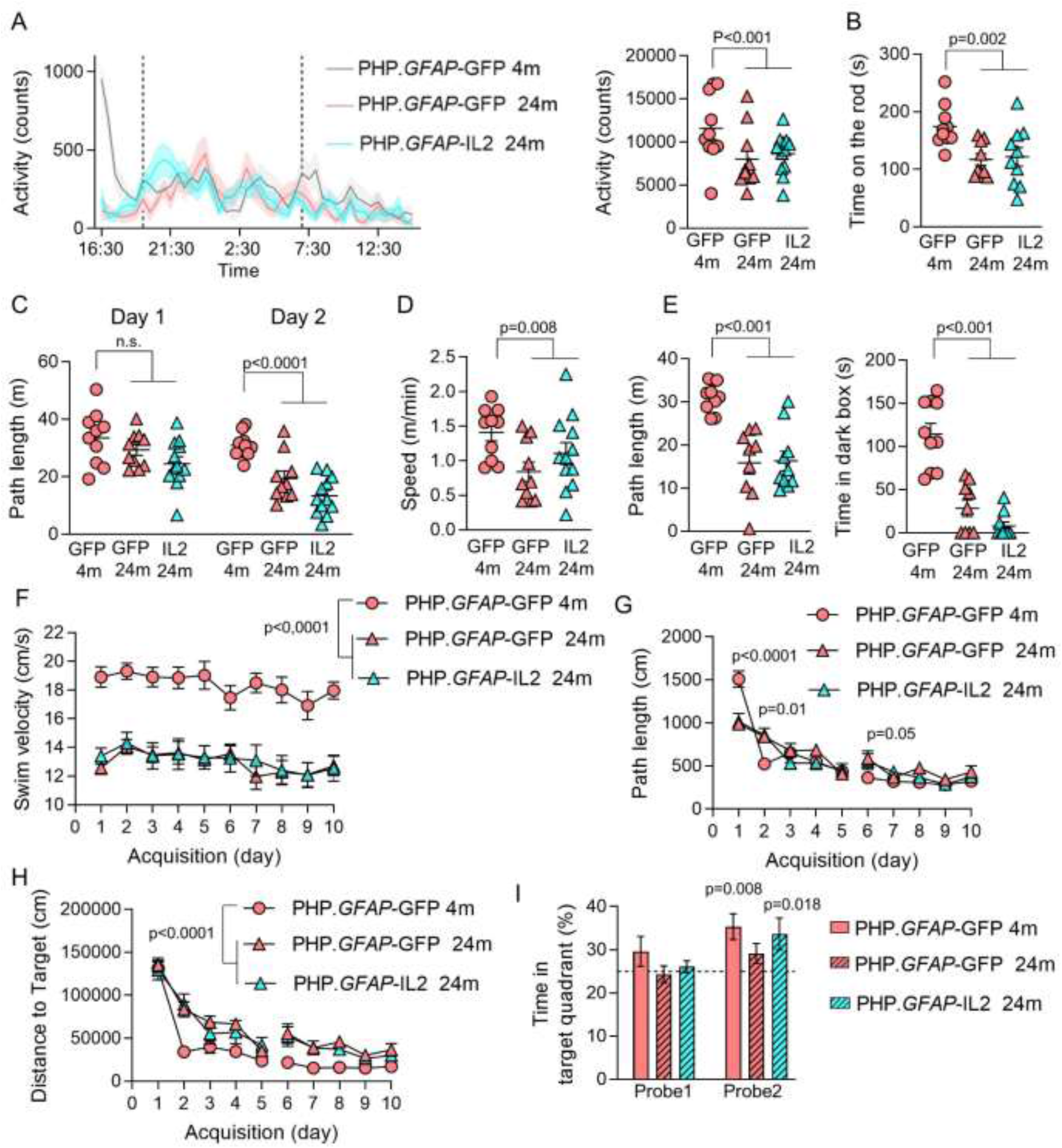
Synthetic IL2 delivery prevents age-induced decline in spatial memory formation without altering age-induced degeneration in mobility or novelty-seeking behaviors. Wildtype mice treated with PHP.*GFAP*-GFP control vector at two months of age were compared to aged mice treated with PHP. *GFAP*-IL2 (or PHP. *GFAP*-GFP control vector) at 22 months of age. Two months post-treatment, behavior was tested. **A)** Home cage activity was assessed through infrared registration of horizontal movements of single housed mice every 30 minutes (n=10,10,12). Left, activity during 24 h. Dotted line indicate day/night boundaries. Right, average activity counts. **B)** Time spent on the rod, average of 4 repeated tests of 300 seconds (n =10,9,10). **C)** Total distance moved in an open field test in trials of 1 day intervals (n=10,10,12). **D)** Exploration during the the sociability test(n=10,10,12). **E)** Light-dark test (n =10,9,10). Left, distance moved in the lighted arena. Right, latency to enter the dark zone. **F)** Spatial learning in the Morris water maze (n = 10,10,12). Swim velocity, **G)** path length to finding the hidden platform, and **H)** distance to target during acquisition trials. **I)** Probe tests after 5 days (probe 1) and 10 days (probe 2) of acquisition in the Morris water maze. Time spent in the Target Quadrant was significantly above chance during the second probe trial in the young control group and in old IL2-treated animals. Mean ±SEM. A and F, 2-way ANOVA repeated measures with age and treatment as the main factors, B-D, 2-way ANOVA with age and treatment as the main factors, G, one-sample t-test to 25% chance level.

## Discussion

The aging of the brain is a multifaceted phenomena, encompassing structural, neuronal, glial and microenvironmental changes. The causative drivers of the cellular changes are difficult to untangle, as alterations such as loss of neuronal circuits and plasticity could be primarily driven by cell intrinsic neuronal-dependent aging or could be derived from the impact of aging glial support. The role of inflammation in this process is likely to be a primary driver of at least some aspects of the aging process, based on correction experiments. For example, defects in learning and memory capacity in the contextual fear conditioning and radial arm water maze assays can be partially corrected in 18 month-old mice through the repeated injection of plasma from young mice [47]. Likewise, heterochronic parabiosis, joining the circulatory system of a 21 month-old mouse to a 2 month-old mouse, improved the performance of old mice in olfactory sensitivity assays [48]. Thus, while a neuronal cell intrinsic aging effect is likely, the emergent feature of cognitive decline must at least partly depend on environmental factors, such as a build up of toxic or inflammatory products. T cells are a key potential player in the development of this neuroaging environment. T cells can directly produce, or stimulate the production of, key inflammatory mediators, including cytokines, ROS and autoantibodies, a process that can directly contribute to emergent features such as the constraint of the neuronal stem cell niche [26]. The identification of T cells as a potential culprit opens the possibility of Tregs as a cure. Tregs have direct antiinflammatory properties, preventing excessively exuberant responses from conventional T cells. In addition, Tregs can produce pro-repair orientated cytokines, such as amphiregulin and osteopontin [39, 49], which are neuroprotective following injuries such as stroke. As Treg numbers in the brain are low [36], it is expected that their influence will be mediated, at least in part, through the reprogramming of glial cells. In the peripheral context, Tregs are capable of reprogramming monocytes into the more pro-repair anti-inflammatory profile [50]. In the brain, an analogous program may be imparted onto microglia [51]. Microglial polarization can, in turn, impact astrocyte polarization [52]. Tregs can also stimulate OPCs and promote myelination, in a pro-repair state [53]. When considering the efficacy of PHP.*GFAP*-IL2 in mitigating aging effects, the parsimonious explanation would be a primary effect via the local expansion of Tregs. When used in the context of traumatic brain injury, the beneficial effects of PHP.*GFAP*-IL2 were only observed in the presence of an adaptive immune system, suggesting that the levels of IL2 achieved (~2pg/ml) are too low to trigger the activation of the lower affinity receptor expressed in non-Treg lineages [43]. While it would be attractive to speculate on the role of individual downstream mediators, the complexity and interdependencies of aging make a multifactorial function more likely, integrating pro-repair, anti-inflammatory and glial reprogramming functions.

Single cell transcriptomics of aging glia identified a shared molecular age-induced signature across all major glial cell types. The pathways identified concord with the current molecular understanding of aging, led at the cellular level by mitochondrial dysfunction, loss of proteostasis and cellular senescence. Mitochrondrial dysfunction (here identified as changes in the oxidative phosphorylation pathway and mitophagy pathway) has been identified as a key cell-intrinsic marker of aging, with dysfunction in microglia a potential contributor to neurodegeneration in dementia [54]. Indeed, an increased burden of mitochrondrial dysfunctional can hasten the progression of brain atrophy in mouse models of AD [55], while mitophagy of defective mitochondria reverses cognitive decline [56]. Loss of proteostasis is another reoccurring theme in aging across tissues, and is consistent here with changes to protein production (RNA transport, spliceosome and ribosome), processing (protein processing in endoplasmic reticulum) and degradation (proteasome) pathways. The mechanistic link between proteostasis and aging is unclear, with a lead contender being the toxic or inhibitory build-up of misfolded and/or damaged proteins. The genetic link between proteostasis network components and neurological diseases, such as ALS, AD and PD, strongly suggest that a failure of normal proteostasis is a distinct risk factor for neuropathology [57]. Related to proteostasis is autophagy, a process intimately linked to aging [58]. AD patients exhibit dysregulated autophagy [59] and loss of autophagy causes severe neurodegeneration in mice [60]. For the specific signaling pathways identified, the Neuregulin-ErbB pathway, Ras pathway and PI3K/AKT/mTOR [61–63] are all integral to neuroaging, and potential drivers for the cellular phenotypes developing in aging glial cells.

Surprisingly, the local provision of IL2 in the brain was sufficient to substantially revert almost all of these age-induced pathway changes across microglia, oligodendrocytes and astrocytes. This was despite relatively few immune pathways being identified as altered, apart from the upregulation of MHCII, previously linked to PHP.*GFAP*-IL2 treatment [43], and alterations in the TLR and IL17 signaling pathways in microglia. Whether this effect is mediated via direct reprogramming, such as Treg cytokine-mediated effects, or indirect effects of microenvironmental cleansing, remains to be seen. It is, however, highly promising that such cell-intrinsic molecular signatures of aging are responsive to late-stage intervention.

While this study suggests that an analog of the PHP.*GFAP*-IL2 treatment could be of use to avert cognitive decline in ageing in humans, there are several key barriers to translation. First, the degree to which the aging process is conserved across the species barrier is unclear. Many of the biological processes occurring in the aged brain, such as accumulation of DNA modifications, mitochondrial dysfunction and loss of proteostasis, are shared across species [64]. Likewise, cognitive decline is common between mice and humans [65], including reduction in spatial navigation performance (such as the Morris water maze in mice, or the equivalent in humans) [66]. Despite this, mouse cognitive decline seems to appear relatively sooner and faster than for humans [67], and decline in elaborate cognitive abilities specific to human, such as language, cannot be tested in animal models. Further, the neurogenic niche inhibited by T cells in aged mice [26] may not exist in humans [68]. Thus, even if the restoration of spatial navigation decline was achieved through brain-specific IL2 delivery in humans, it is not clear that other aspects of cognitive decline, such as language deficits, share a common cellular or molecular cause, or would respond to the same treatment approach. A second potential limitation to translation is the kinetics involved in cognitive decline. Here we treated aged mice for two months prior to assessment, with the treated aged mice performing similar in spatial navigation capacity to young control mouse. It remains unknown, however, whether the treatment actively reversed the molecular, cellular and behavioral processes of aging, or whether it merely prevented the decline. The kinetics of cognitive decline are compressed into a mouse lifespan, raising the question of whether treatment in humans could impact a decline occurring over the course of decades. If cognitive decline is primarily driven by active changes in the brain microenvironment, such as increased basal inflammation induced by a lifelong stimulation of the immune system [18, 19], then even transient pulses of purification could reverse or prevent the decline. On the other hand, if cognitive decline is driven primarily by programmed senescence, then such a treatment could be expected to, at most, delay cognitive decline. Fortunately, experiments in mice largely support the former model over the latter, as murine cognitive decline occurs with age even though individual mouse neurons can long out-survive a mouse, when transplanted into a longer-lived host [69]. Finally, the molecular pathways of the treatment itself need to be considered. The delivery system used here, the PHP.B capsid, performs poorer in blood-brain-barrier crossing in non-human primates than it does in mice [70]. Fortunately, alternative AAV capsids are available that perform well in humans [71], and sustained cargo expression in the brain has been observed to span years [72], an advantage for a potential longevity treatment. Tregs are found in both the mouse and human brain [36], and the IL2 pathway is highly conserved across the species, so it is likely that the expansion of brain Tregs could be induced in humans if cargo delivery was achieved. Nonetheless, substantial technical and regulatory barriers exit when considering the development of any longevity treatments that are designed for the use of healthy individuals, with long-term safety data being essential.

## Supporting information

Supplementary Resource 2

Supplementary Resource 1

## Acknowledgements

The work was supported by the VIB, an ERC Consolidator Grant TissueTreg (to A.L.), an ERC Proof of Concept Grant TreatBrainDamage (to A.L.), FWO Research Grant 1503420N (to EP), an SAO-FRA pilot grant (20190032, to E.P.), an ERC Starting Grant AstroFunc (to M.G.H.), ERC Proof of Concept Grant AD-VIP (to M.G.H.), ERNAET Chair NCBio (to MGH), and the Biotechnology and Biological Sciences Research Council through Institute Strategic Program Grant funding BBS/E/B/000C0427 and BBS/E/B/000C0428, and the Biotechnology and Biological Sciences Research Council Core Capability Grant to the Babraham Institute. E.P. was supported by a fellowship from the FWO. The authors acknowledge the important contributions of Jeason Haughton (VIB) for mouse husbandry, Pier-Andrée Penttila and the KUL FACS Core, and the VIB Single Cell Sequencing Core. The VIB and Babraham Institute have filed a patent application covering aspects of the work included in the publication and are pursuing commercialisation.

