## Supplementary methods and figures for "Molecular and cognitive signatures of ageing partially restored through synthetic delivery of IL2 to the brain"

### Materials and methods

*Mice*

C57BL/6 mice at 2 and 22 months of age were purchased from Janvier Labs and housed in SPF conditions, under a 12 hour light/dark cycle in a temperature and humidity-controlled room with *ad libitum* access to food and water. All animal procedures were approved by the KU Leuven Animal Ethics Committee (P124/2019), taking into account relevant national and European guidelines. Male mice were used in this study, unless otherwise specified. Sample sizes for mouse experiments were chosen based on power calculations and pilot data, in conjunction with the Animal Ethics Committee, to allow for robust sensitivity without excessive animal use. Mice were selected randomly for inclusion into the various experimental groups, with the animal technicians performing experimental procedures and clinical measurements blinded as to the identity of experimental groups.

*Behavioral experiments*

Behavioral experiments were performed in 4 months and 24 months old males and compared to littermate controls. Mice were habituated to their new environment for at least 5 days and tests were conducted during the light phase of their activity cycle. Tests were performed and analyzed by an observer blind to the genotype or treatment group. If tracking error(s) occurred, mice were excluded.

**Cage Activity**. Ambulatory behavior was investigated over a 23 h time-period, starting at 4 p.m. until 3:30 p.m. the following day [1]. Lights were switched off at 7pm, and on again at 7am. During this test, animals were individually housed in transparent cages (20 × 26 cm) with minimal bedding, chow and water and placed in a laboratory-built activity logger with three infrared beams. Infrared beams register horizontal movements; beam breaks were recorded over 30 min time bins.

**Open Field**. Open-field exploration was tested in a 50 cm × 50 cm × 30 cm (w×l×h) square arena illuminated by indirect light. Animals were dark adapted for 30 min and tested in the arena for 10 min. Movements of the mice were video-tracked for 10 minutes using ANY-maze Video Tracking System software (Stoelting Europe).

**Light/dark test.** The apparatus used for the light/dark test consisted of a cage (50 × 50 × 30 cm) divided into two compartments by a partition with a door. One compartment was brightly illuminated, while the other was dark. Mice were placed in the light compartment and allowed to move freely between the two chambers for 10 min. Mouse activity in the light compartment was tracked using ANY-maze Video Tracking System software (Stoelting Europe).

**Rotarod test.** Motor coordination and equilibrium were tested with a rotarod setup (MED Associates Inc.). Mice were first trained at constant speed (4 rpm, 2 min) and then tested on four trials (intertrial interval, 10 min). During the test trials, the animals had to balance on a rotating rod that accelerated from 4 rpm to 40 rpm in 300 seconds. Latency to fall off the rod was recorded, up to a 5 min cut-off point.

**Morris Water Maze (MWM).** Spatial learning and cognitive flexibility were tested in the hidden platform Morris water maze (MWM). A circular pool (150 cm diameter) was filled with opacified (0.01% Acusol OP301, Dow Chemicals) water (26 ± 1°C). The platform (15 cm diameter) was hidden 1 cm underneath the surface of the water. For spatial learning, the mice were trained for 10 days to a fixed platform position [2, 3]. To evaluate reference memory, probe trials (100 seconds) were conducted on days 6 and 11 during acquisition learning. During probe trials, floater mice were excluded. The escape platform was removed from the pool and mice were allowed to explore the maze for 100 seconds. Swim paths were tracked with Ethovision software (Noldus).

**Sociability test.** Sociability was evaluated using the three-chamber test. The set-up consists of a rectangular transparent Plexiglas box divided into three compartments, separated by two partitions. Multiple holes in the partitions allow sniffing interactions between a mouse in the central chamber (42 x 26 cm) and a mouse in either the left or right chamber (26 x 26 cm). The test consisted of two consecutive stages: acclimation stage and a sociability stage. After habituation to the central compartment (5 min), a stranger mouse (same sex) was placed in one of the side chambers while the other was left empty. Approach behavior to side compartments was recorded for 10 min (sociability). Animal behavior was recorded using a webcam and ANY-maze Video Tracking System software (Stoelting Europe).

*Statistics*

Comparisons between two groups were performed using unpaired two-tailed Student's t tests. *Post hoc* Holm's or Dunnett's multiple comparisons tests were performed, when required. Two-way ANOVA was used when appropriate, with age and treatment as the main factors. Cage activity behavior and MWM acquisition was analyzed with 2-way ANOVA with repeated measures, with age and treatment as the main factors. Non-parametric testing was performed when data was not normally distributed (QQ plot for visual check and Shapiro-Wilk normality test on pooled residuals). The value of n reported within figure legends represents the number of animals, unless otherwise specified. Values are represented as mean ± SEM.

*AAV vector production and purification*

AAV-PHP.B production was performed by Vigene Sciences (Rockville, MD, USA) or VectorBuilder (Neu-Isenburg, Germany), using the classical tri-transfection method, with subsequent vector titration performed using a qPCR-based methodology [4, 5]. For AAV-PHP.B.GFAP-IL2, the mouse IL2 coding sequence, together with 5' and 3' UTR (accession number BC116845) was cloned into a single stranded AAV2-derived expression cassette, containing a full-length GFAP promoter [6], woodchuck hepatitis post-transcriptional regulatory element (WPRE) and bovine growth hormone polyadenylation (bGH polyA) sequence. Control vectors were prepared by swapping the IL2 coding sequence for that encoding enhanced green fluorescent protein (EGFP, Vector Biolabs). The vector (100 µl total volume) was administered to mice via the lateral tail vein at $1x10^9$ vector genomes/dose. Mice were used for experimental procedures 2 months after AAV injection, unless otherwise indicated.

*Single-cell RNA sequencing*

Mice were deeply anaesthetized with an intraperitoneal injection of a ketamine (87 mg/kg), xylazine (13 mg/kg) mixture and transcardially perfused with ice cold PBS. The brains were put into brain medium: DMEM (Gibco) supplemented with B27 (Thermo Fisher), 2 mM Sodium pyruvate (Gibco), 500 µM N-acetyl-cysteine (Sigma Aldrich) and 5 uM Actinomycin D (Sigma Aldrich). The brains were then cut into small pieces and digested in brain medium with 33 UI/mL papain (Sigma Aldrich) and 40 µg/mL DNase I (Sigma Aldrich) for 30 minutes at 37ºC. Digested tissue was mechanically disrupted, filtered through 100 µm mesh and enriched by density centrifugation (300 $g_{Av}$, 11 minutes, no brake) through 24% Percoll (GE Healthcare). Cell sorting was performed using a Sony MA900 with a 100 µm nozzle, using an antibody panel including eBioscience™ Fixable Viability Dye eFluor™ 780, CD11b, CD45, CX3CR1 (for microglia), O4 (for oligodendrocytes), ACSA-2 (for astrocytes) and PDGFRα (for OPCs) for sorting. Sorted cells were mixed per mouse and resuspended in DMEM with 10% FBS. Post-sort, the cell count and the viability of the samples were confirmed using a LUNA-FL dual fluorescence cell counter (Logos Biosystems). Post-cell count and quality control, the samples were immediately loaded onto the Chromium Controller. For each experiment, approximately 8,700 cells were added to each channel for a targeted cell recovery of 5,000 cells. Library preparations were performed using the 10X Genomics Chromium Single Cell 3' Kit, v3 (10X Genomics). Libraries were prepared according to manufacturer's instructions (Single cell 3' reagent kits v3 user guide; CG000183 Rev C), and at the different recommended check points the library quality was assessed using a Qubit 2 Fluorometer (ThermoFisher) and a Bioanalyzer HS DNA kit (Agilent). With a sequencing coverage targeted for 50,000 reads per cell, single cell libraries were sequenced on an Illumina Novaseq 6000 or Illumina HiSeq platform using a paired-end sequencing workflow with recommended 10X, v3 read parameters (28-8-0-91 cycles).

Fastq from the raw sequencing data underwent quality control (QC) analysis and were converted to counts data using 10X Genomics CellRanger v4.0.0 counts pipeline [7]. QC parameters were found to be within the expected ranges for single cell sequencing data, as per 10X Genomics user guides. The counts data was further analyzed using Seurat v4.0.2 [8] in R v4.0.5 [9]. The data was filtered to remove cells low quality cells, classed as those containing >10% mitochondrial genes [10]. The individual counts data was then combined and normalized by the variance stabilizing transformation method, using the sctransform v0.3.2 library [11], correcting for batch and regressing for RNA counts and percentage mitochondrial gene expression. The normalized data was then analyzed for detection and removal of multiplets using scDblFinder v1.4.0 [12].

Population markers and differential expression was performed using the built-in methods in Seurat, using the negative binomial test. Likewise, PCA and UMAP projections were also performed using Seurat. External data was downloaded directly from the Mouse Brain Atlas (mousebrain.org/downloads; [13]) for mapping onto the astrocyte data generated in this study, using the data integration pipeline in Seurat. Pseudotime analyses were performed and trajectories were constructed using the DDRTree algorithm in Monocle v2.22.0 [14]. Gene set enrichment and pathway enrichment were performed using GAGE v2.44.0 [15] and Pathview v1.34.0 [16], respectively. The full analysis code is available on GitHub (https://github.com/AdrianListon/AAV-IL2) with the data available on GEO as dataset GSE190486.

**Supplementary Resource 1. Differential expression with age and treatment across glial cell types.** Wildtype mice were treated with PHP.*GFAP*-IL2 (or PHP.*GFAP*-GFP control vector) at 4 months or 24 months of age (n=3/group). Two months post-treatment, the glial compartment was sorted from perfused mice and assessed via 10x single-cell transcriptomics. For the microglia, oligodendrocyte and astrocyte clusters, differential expression analysis was performed. Differentially-expressed genes are listed for age and treatment, together with p value, adjusted value, average $\log_2$ fold-change and the fraction of cells expressing the gene in group 1 (first listed sample) and group 2 (second listed sample).

**Supplementary Resource 2. Differential pathway analysis with age and treatment across glial cell types.** Gene set enrichment was performed based on differential expression across various comparisons, including age (aged PHP.*GFAP*-GFP mice versus young PHP.*GFAP*-GFP mice), treatment in young mice (young PHP.*GFAP*-GFP mice versus young PHP.*GFAP*-IL2 mice), treatment in old mice (aged PHP.*GFAP*-IL2 mice versus aged PHP.*GFAP*-GFP mice) and age in treated mice (aged PHP.*GFAP*-IL2 mice versus young PHP.*GFAP*-IL2 mice). Gene sets were taken from the GAGE library and curated to the respective KEGG pathway. For each gene set enriched in at least one of the four comparisons, the stat mean and q value for each comparison is shown. The final column indicates the number of comparisons where the q value reached the threshold for statistical significance.

**Supplementary Figure 1. Identification of major glial populations based on key transcriptional markers.** Young and old wildtype mice, treated with PHP.*GFAP*-IL2 (or PHP.*GFAP*-GFP control vector) were assessed two months post-treatment by glial single cell sequencing using 10x single-cell transcriptomics. UMAP feature expression plots for key microglial markers (*Tmem119, P2ry12, Cx3cr1, Hexb*), oligodendrocyte markers (*Mog, Cldn11, Mobp, Mbp*), astrocyte markers (*Aqp4, Fgfr3, Slc4a4, Sox9*) and oligodendrocyte precursor cell (OPC) markers (*Pdgfra, C1al1, Olig2, Neu4*).

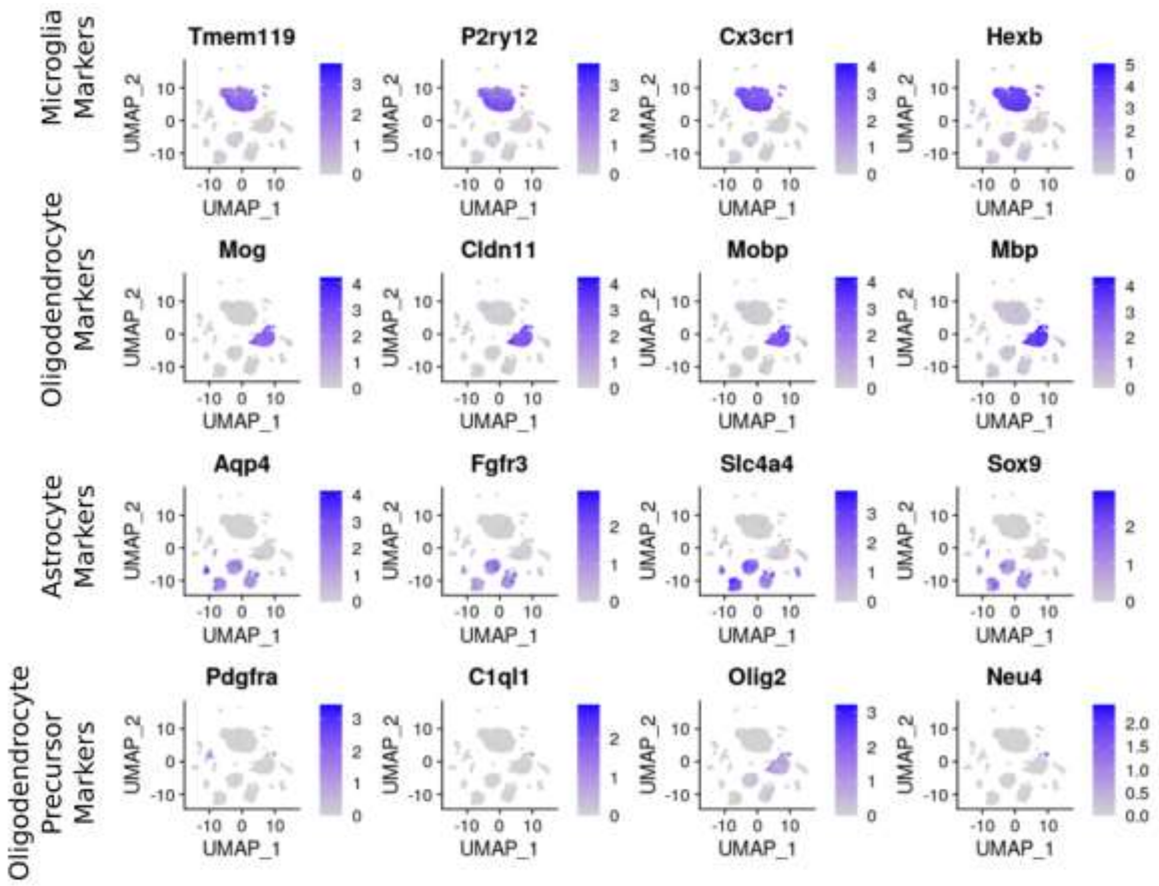

**Supplementary Figure 2. Identification of non-target populations based on key transcriptional markers.** Young and old wildtype mice, treated with PHP.*GFAP*-IL2 (or PHP.*GFAP*-GFP control vector) were assessed two months post-treatment by glial single cell sequencing using 10x single-cell transcriptomics. UMAP feature expression plots for markers characteristic of non-target populations, aiding the annotation of cellular origin of contaminates into **A)** reticulocytes, **B)** neutrophils, **C)** perivascular macrophages, **D)** Choroid plexus epithelial cells, **E)** neurons, **F)** olfactory ensheathing cells, **G)** olfactory neurons, **H)** interneurons, **J)** neuro stem and progenitor cells, **K)** ependymal cells, **L)** hypendymal cells, **M)** vascular endothelial cells, **N)** vascular leptomeningeal cells, **O)** pericytes and **P)** vascular smooth muscle cells.

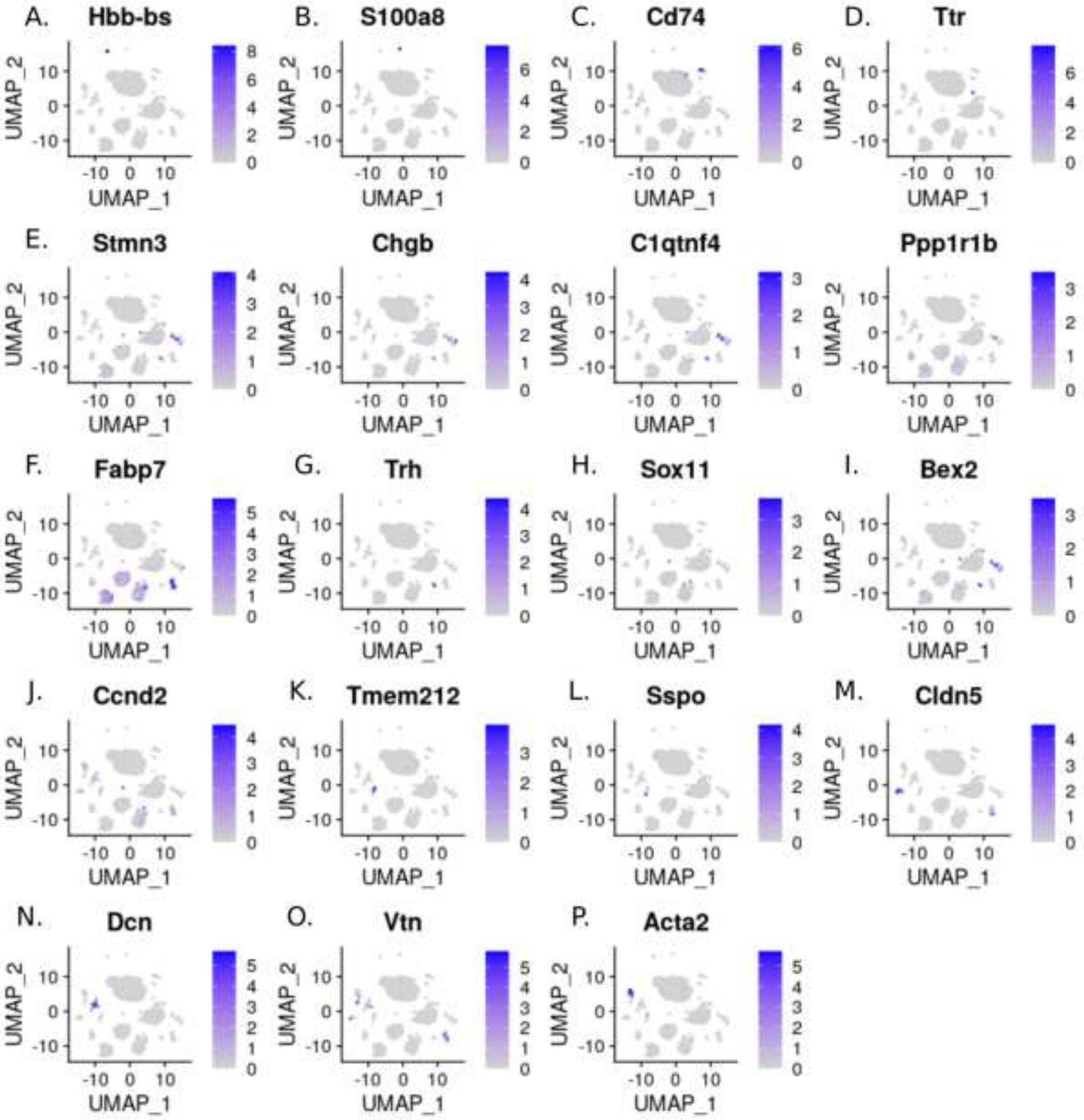

**Supplementary Figure 3. Expression of key homeostatic and activation markers within the astrocytic populations.** Young and old wildtype mice, treated with PHP.*GFAP*-IL2 (or PHP.*GFAP*-GFP control vector) were assessed two months post-treatment by glial single cell sequencing using 10x single-cell transcriptomics. Astrocytes were identified based on marker expression and reclustered/reprojected via UMAP. **A)** UMAP feature expression plots for Bergmann glia, **B)** cerebellar astrocytes, **C)** non-telencephalon astrocytes, **D)** olfactory astrocytes, **E)** striatal astrocytes and **F)** telencephalon astrocytes. **G)** Astrocyte expression data was mapped onto the astrocyte expression data available at mousebrain.org, using the Seurat integration pipeline to verify identified populations.

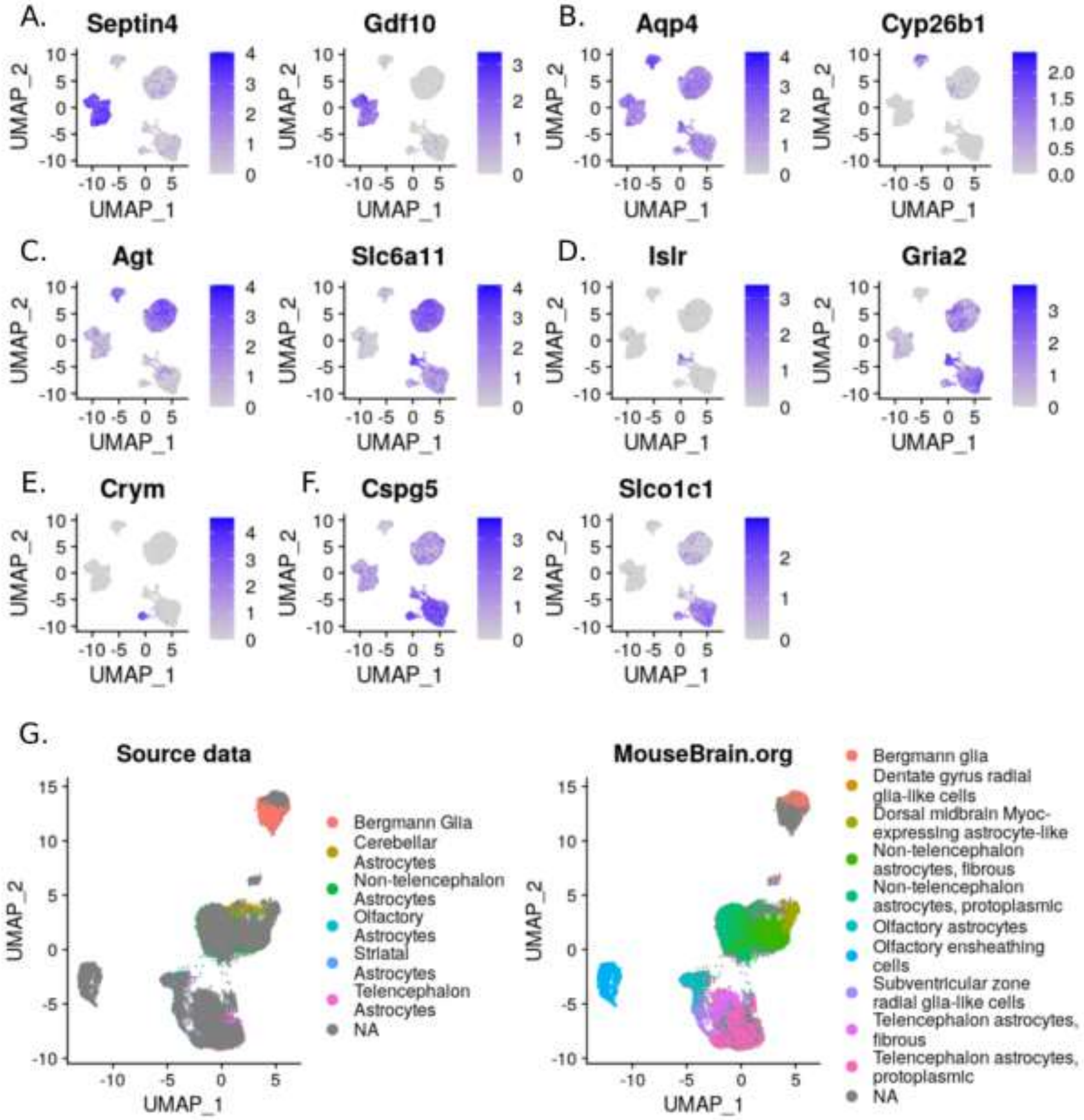

**Supplementary Figure 4. Expression of key homeostatic and activation markers within the microglial populations.** Young and old wildtype mice, treated with PHP.*GFAP*-IL2 (or PHP.*GFAP*-GFP control vector) were assessed two months post-treatment by glial single cell sequencing via 10x single-cell transcriptomics. Microglia were identified based on marker expression and reclustered/reprojected via UMAP. UMAP feature expression plots for **A)** homeostatic markers (*Cx3cr1*, *P2ry12*, *Tmem119*, *Cst3*, *Hexb*) and **B)** activation-associated markers (*Apoe*, *Lpl*, *Cst7*, *Axl*, *Itgax*, *Spp1*, *Ccl6*, *Csf1*).

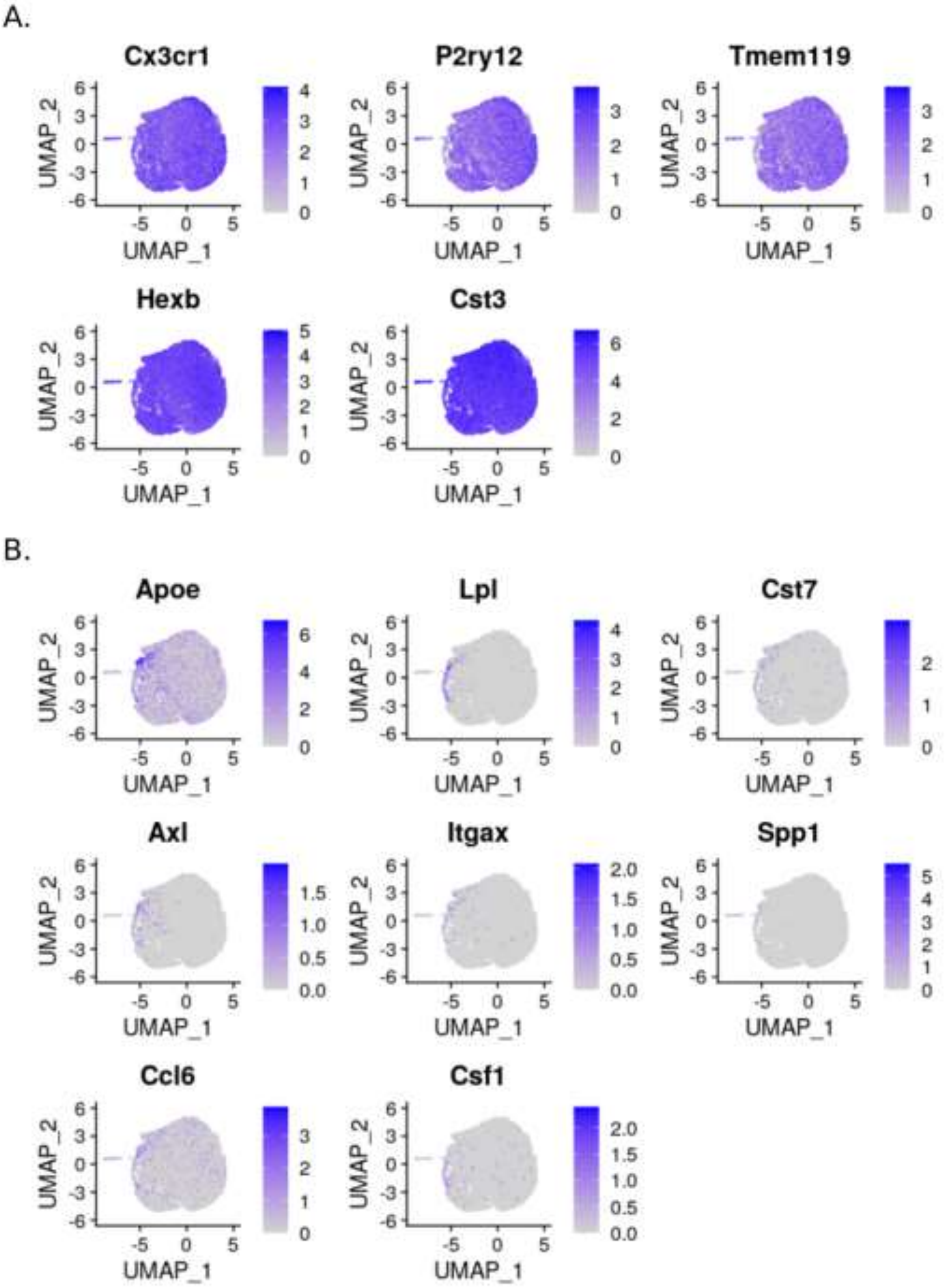

**Supplementary Figure 5. Normal pseudotime cellular trajectory progression for oligodendrocytes following IL2-treatment.** Wildtype mice were treated with PHP.*GFAP*-IL2 (or PHP.*GFAP*-GFP control vector) at 2 months or 22 months of age (n=3/group). Two months post-treatment, the glial compartment was sorted from perfused mice and assessed using 10x single-cell transcriptomics. Oligodendrocytes and OPC clusters were reclustered and a pseudotime trajectory constructed (branching trajectory trees), using the DDRTree algorithm in Monocle v2. The tree was rooted at the branch with the highest proportion of young cells treated with PHP.*GFAP*-GFP. The trees were generated over the whole data set and illustrated separately for PHP.*GFAP*-GFP and PHP.*GFAP*-IL2.

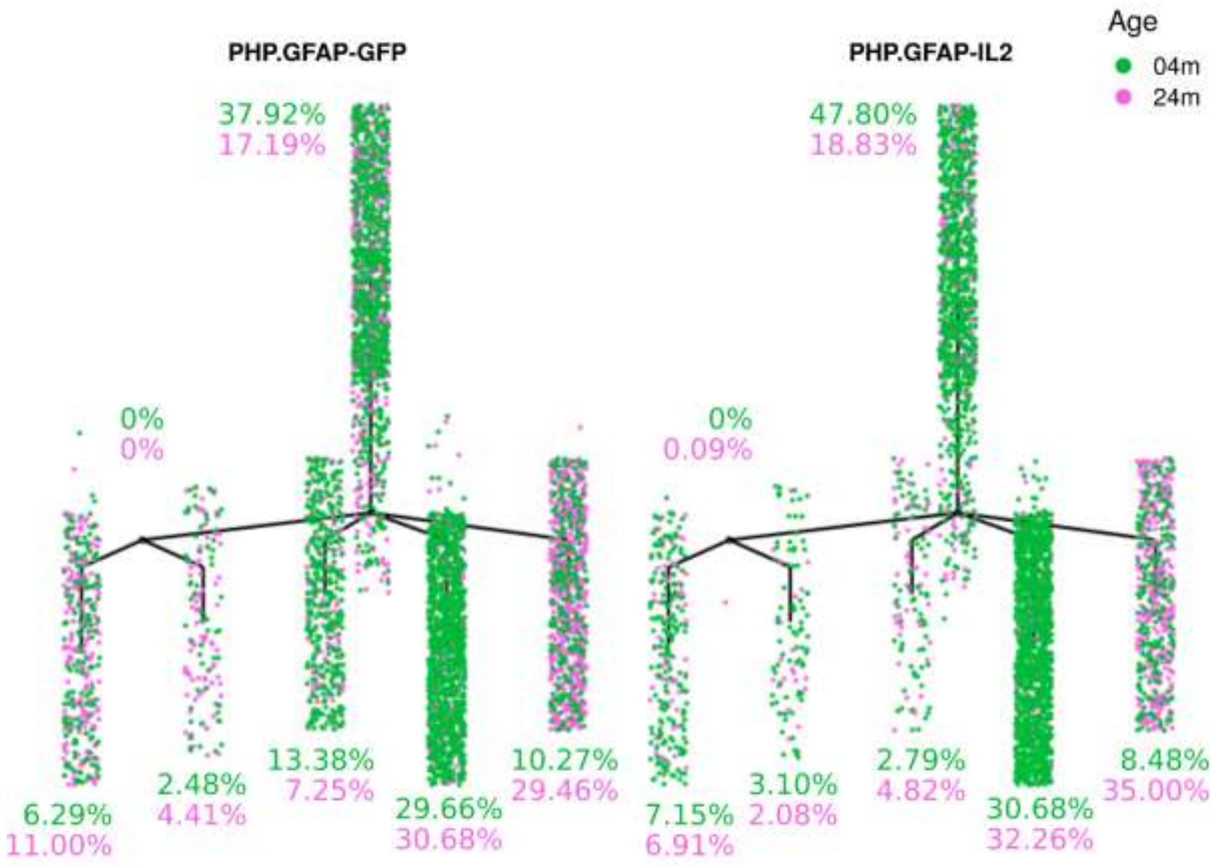

**Supplementary Figure 6. Expression of key homeostatic and activation markers within the oligodendrocyte and OPC populations.** Young and old wildtype mice, treated with PHP.*GFAP*-IL2 (or PHP.*GFAP*-GFP control vector) were assessed two months post-treatment using 10x single-cell transcriptomics. Oligodendrocytes (Oligodendrocyte subgroups 1-5; Oligos 1-5) and oligodendrocyte precursor cells (OPCs) were identified based on marker expression and reclustered/reprojected into UMAP space. UMAP feature expression plots for key markers enriched in **A)** OLIGO1, **B)** OLIGO2, **C)** OLIGO3, **D)** OLIGO4, **E)** OLIGO5, and **F)** Oligodendrocyte precursor clusters.

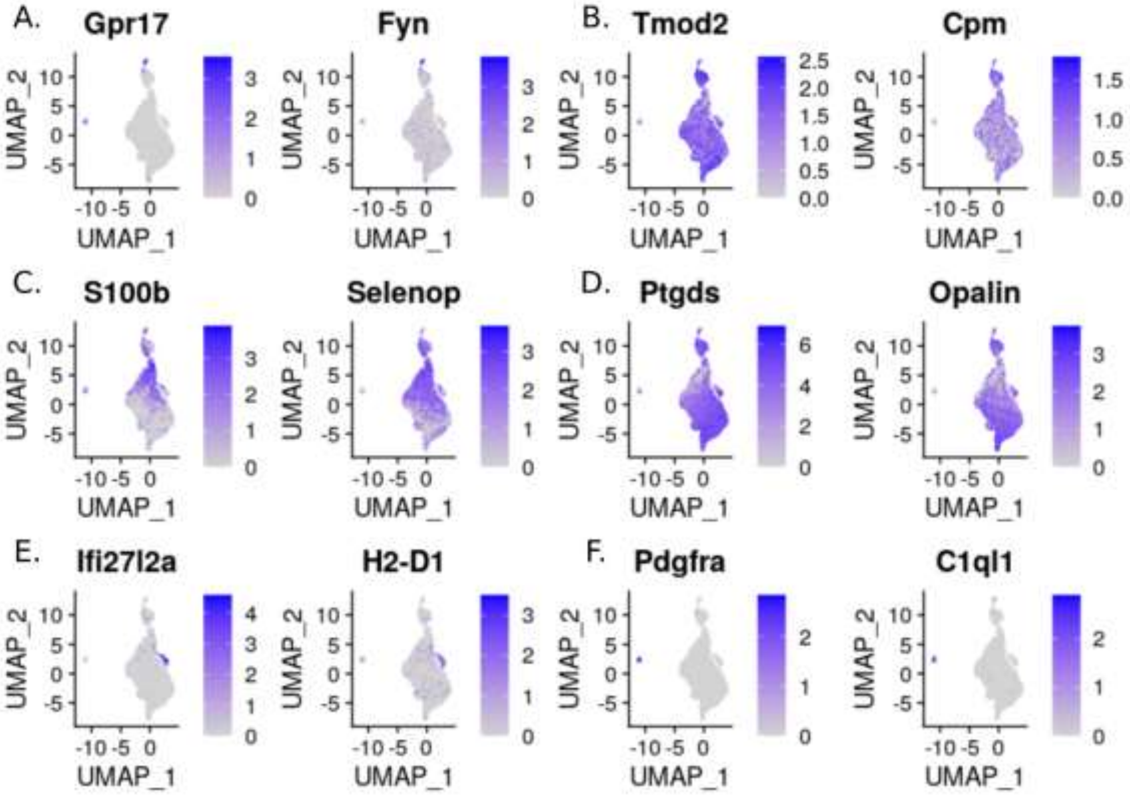

**Supplementary Figure 7. Pseudotime trajectory trees for astrocytes during age and treatment with IL2.** Young and old wildtype mice, treated with PHP.*GFAP*-IL2 (or PHP.*GFAP*-GFP control vector) were assessed two months post-treatment by single cell sequencing using 10x single-cell transcriptomics. Astrocytes were identified based on marker expression and reclustered/reprojected via UMAP. Astrocyte subclusters, annotated based on key markers, were assessed for pseudotime trajectory. Pseudotime analysis generated branching trajectory trees of cells based on the gene expression profile of each cell, using the DDRTree algorithm in Monocle v2. The tree was rooted at the branch with the highest proportion of young cells treated with PHP.*GFAP*-GFP. The trees were generated over the whole data and illustrated separately for PHP.*GFAP*-GFP and PHP.*GFAP*-IL2 for **A)** Bergmann glia, **B)** cerebellar astrocytes, **C)** non-telencephalon astrocytes, **D)** olfactory astrocytes, **E)** striatal astrocytes and **F)** telencephalon astrocytes.

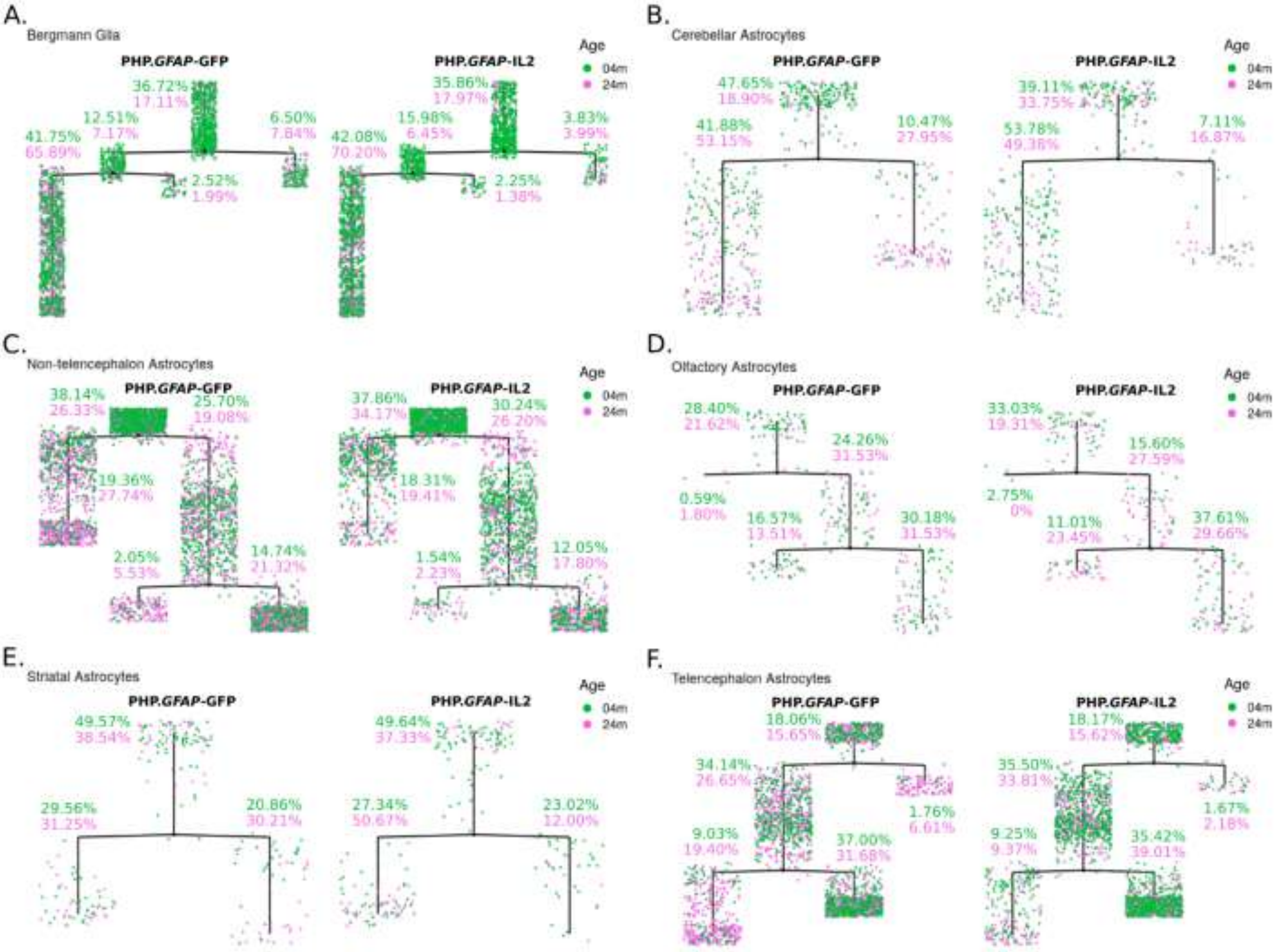
